# Sex-specific vulnerability to behavioural and morphological alterations in a mouse model of Parkinson’s disease overexpressing mutant A53T alpha-synuclein

**DOI:** 10.1101/2025.07.30.667616

**Authors:** M Zubelzu, R Bidgood, A Murueta-Goyena, JA Ruiz-Ortega, JV Lafuente, T Morera-Herreras

**Author notes:** Corresponding author: **Raphaelle Bidgood**, Department of Pharmacology, Faculty of Medicine and Nursing, University of the Basque Country (EHU), E-mail address, Postal address: Department of Pharmacology, Faculty of Medicine and Nursing, University of the Basque Country (EHU), Barrio Sarriena s/n, 48940-Leioa, Spain.

## Abstract

Parkinson’s disease (PD) is characterised by progressive degeneration of dopaminergic neurons in the *substantia nigra pars compacta* (SNc) and accumulation of misfolded α-synuclein (α-syn). Neuroinflammation also contributes to disease onset and progression. Notably, PD exhibits sexual dimorphism in clinical presentation and treatment response. This study investigated sex-specific behavioural and morphological changes in a mouse model overexpressing A53T α-syn.

Male and female C57BL/6J mice received bilateral intranigral injections of adeno-associated viral vectors encoding mutant A53T α-syn or empty vectors. Motor function was assessed at 60 and 120 days post-surgery using open field, wire hang, pole, and balance beam tests. Brains were collected for immunohistochemical analyses of α-syn pathology, nigrostriatal integrity (tyrosine hydroxylase, TH), axonal degeneration, and neuroinflammation.

α-Syn overexpression induced early, subtle motor deficits primarily in males, despite preserved SNc neuronal density. Automated analysis of balance beam walking behaviour (DeepLabCut, SimBA) revealed increased immobility and reduced walking time in α-syn males. At 120 days only, striatal TH levels were significantly reduced, driven by reductions in α-syn males. Although undetected at 60 days, an axonal degeneration index (combining striatal TH optical density and axonal swellings) revealed more advanced degeneration in α-syn males, suggesting faster disease progression. At both time points, α-syn mice showed increased striatal astrogliosis without sex differences, indicating α-syn-associated neuroinflammation.

These findings support a PD model of early axonal degeneration and reactive astrogliosis preceding neuronal loss. The sex-specific behavioural and neuropathological patterns underscore the importance of incorporating sex as a biological variable in preclinical models and developing tailored therapeutic strategies.

## 1. Introduction

Parkinson’s disease (PD) is a progressive neurodegenerative disorder characterised primarily by motor dysfunction, including bradykinesia, rigidity, resting tremor, and postural instability, as well as a range of non-motor symptoms (Mirelman et al., 2019; Tysnes and Storstein, 2017). The pathological hallmark of PD is the progressive loss of dopaminergic neurons in the *substantia nigra pars compacta* (SNc), leading to striatal dopamine depletion. In addition, PD is marked by the accumulation of misfolded α-synuclein (α-syn) protein aggregates, forming Lewy bodies and Lewy neurites, which are considered central to disease pathogenesis (Blesa et al., 2022; Choong and Mochizuki, 2022; Sharma and Burré, 2023).

Observational clinical studies have consistently reported sex differences in PD incidence, progression, and symptomatology. Men are more likely to develop PD than women (Weber and Clyne, 2021), but women often experience a higher burden of non-motor symptoms such as anxiety, depression, and dysautonomia (Zachry et al., 2021). The biological underpinnings of these sex differences remain poorly understood but are thought to involve complex interactions between sex hormones, genetic factors, and neuroinflammatory responses (Rugbjerg et al., 2013; Zachry et al., 2021).

α-Syn plays a central role in PD pathogenesis, both as a principal component of Lewy pathology and as a driver of neuronal dysfunction and degeneration. Overexpression of mutant forms of α-syn, such as the A53T variant linked to familial PD, recapitulates key pathological features of the disease (Sharma and Burré, 2023). Viral vector-mediated overexpression of α-syn using adeno-associated viruses (AAV) has become a widely used experimental approach to model PD in rodents, allowing targeted, controlled expression of α-syn in specific brain regions (Koprich et al., 2017). These models successfully induce dopaminergic neurodegeneration, motor deficits, synucleinopathy, and neuroinflammation, providing valuable platforms for studying disease mechanisms and testing therapeutic interventions.

Despite increasing use of AAV-mediated α-syn overexpression models, most studies have not systematically examined potential sex differences in α-syn-driven pathology and behavioural outcomes. This represents a significant gap in the current knowledge, particularly given the sex differences observed in human PD. Understanding how sex modulates vulnerability to α-syn-induced neurodegeneration could provide critical insights into disease mechanisms and contribute to the development of more effective, personalised therapeutic strategies.

In the present study, we aimed to investigate whether AAV-mediated overexpression of A53T α-syn induces sex-specific differences in behavioural, neurochemical, and neuropathological outcomes in mice. We systematically assessed motor behaviour, dopaminergic neurodegeneration, and neuroinflammatory responses in both male and female animals over time, to better understand how sex influences the progression of α-syn-mediated pathology.

## 2. Material and methods

### 2.1. Animals and ethical statement

7-week-old male and female C57BL/6J mice (Janvier Labs) were housed in groups of 3-6 in individually ventilated cages under standard laboratory conditions (22±1°C, 55±5 % relative humidity, and a 12:12 h light/dark cycle, with the light phase from 8am to 8pm) with food and water provided *ad libitum*. Every effort was made to minimise animal suffering and to use the minimum number of animals per group and experiment. The experimental protocol was reviewed and approved by the Local Ethical Committee for Animal Research of the University of the Basque Country (EHU, CEEA, ref. M20/2019/219, M20-2022-215 and M30/2019/220). All experiments were performed in accordance with the European Community Council Directive on “The Protection of Animals Used for Scientific Purposes” (2010/63/EU) and with the Spanish law (RD 53/2013) on the care and use of laboratory animals.

### 2.2. Experimental design

This study used a total of 28 male (21–25 g) and 27 female (16–22 g) C57BL/6J mice. AAV vectors encoding for mutant A53T human α-syn or empty vectors were injected bilaterally into the SNc. One female mouse that had been injected with the mutant A53T α-syn was excluded due to the incorrect viral diffusion in one hemisphere.

To investigate the behavioural and morphological alterations in neurons and glial cells within the nigrostriatal pathway, the animals were sacrificed at 60 or 120 days after the stereotaxic injection.

Two batches of animals were used in this study: 16 mice in batch 1 and 40 mice in batch 2. The animals of batch 1 were sacrificed at 60 days post-surgery and those of batch 2, at 120 days. The mice of batch 2 also underwent behavioural testing at 60 days. Immunohistochemistry was carried out independently at both time points. The sample sizes for each study are detailed in Table S1.

### 2.3. AAV surgical procedure

AAV vectors (rAAV9-CMVie/SynP-WPRE) encoding either mutant A53T human α-syn (1×10^13^ genomic particles/ml) or the empty vector (3.1×10^12^ genomic particles/ml) were obtained from the University of Bordeaux (France). Mice were anaesthetised with isoflurane (induction 5%; maintenance 1.5-2%) in oxygen-enriched air (1-2%) and placed in a stereotaxic frame (David Kopf® Instruments). Body temperature was maintained at ∼37°C throughout the experiment using a heating pad connected to a rectal probe. AAV vectors were then bilaterally injected into the SNc (1 μl per hemisphere) using a glass pipette (coordinates from Bregma: anteroposterior (AP) –2.9 mm; mediolateral (ML) ±1.4 mm and dorsoventral (DV) –4.5 mm). The pipette was left in place for 5 minutes after the injection to prevent leakage. Meloxicam (2 mg/kg) was administered subcutaneously to relieve pain and/or reduce inflammation.

### 2.4. Behavioural assessment

Several behavioural tests were performed at 60 and 120 days post-surgery to investigate the possible motor changes induced by the overexpression of mutant A53T human α-syn (Fig. S1). These behavioural tests are explained briefly below. For a more detailed explanation, please refer to the Supplementary Information.

#### 2.4.1. Open field test

The open field test is a common measure of spontaneous motor activity in mice. In this test, a mouse was placed in the center of a square arena, and its movements were automatically recorded using Actitrack software. The arena uses infrared beams to track the animal’s motion accurately. Mice were given 10 minutes to explore (acclimatisation), followed by 10 minutes of data collection. The system measured various parameters, including total activity, total distance travelled, resting time, vertical activity, speed and time spent performing fast, slow and stereotyped movements. The automated tracking also distinguished between central and peripheral zones of the arena, allowing the analysis of the time the mouse spent in each area.

#### 2.4.2. Wire hang test

The wire hang test (grid test) was used to evaluate motor coordination and muscle strength in mice. Each mouse was placed onto a standard cage grid, which was gently shaken to encourage gripping, and then turned upside down approximately 50 cm above a sawdust-filled cage. The time it took for each mouse to fall was recorded, with a maximum test duration of 15 minutes. The test was conducted on two consecutive days.

#### 2.4.3. Balance beam test

The beam-walking test is used in rodents to measure subtle motor and balance impairments that may not be detected by other tests like the rotarod. Mice were trained and then tested on their ability to walk across a narrow (1 cm wide), elevated (60 cm high) 80 cm-long wooden beam (Fig. S2A). Each mouse performed three trials, separated by 10 minutes, and mouse performances were then averaged. Trials were recorded using two cameras: one for manual quantification of hind paw slips (Fig. S2B) and another for automated tracking of the animal to analyse mouse walking pattern along the beam. The automated analysis pipeline used DeepLabCut (Nath et al., 2019) for pose estimation (tracking 5 keypoints) and Simple Behavioral Analysis (SimBA, Goodwin et al., 2024) for behavioural classification. The beam was divided into 10 regions of interest (ROIs), each 10 cm long, to measure the time to cross the beam and the distance travelled. If a mouse never crossed the start line, the trial was excluded from the automated analysis and given the default values of a failed trial (60 s and 0 cm).

#### 2.4.4. Pole test

The pole test was used to assess motor dysfunction in mice. Each mouse was placed facing upwards to the top of a 55 cm-long, 1 cm-diameter pole (Fig. S3). Parameters such as the latency to turn downwards (time the mouse takes to make a 180° turn), the time to descend, and the total time spent on the pole were measured manually. The test was carried out over two consecutive days: three training trials on the first day and three final trials on the second day. The final trials were recorded with a front-view camera for subsequent manual analysis, and mouse performances were then averaged.

### 2.5. Immunoassays

#### 2.5.1. Tissue Processing

Animals were deeply anesthetised and perfused transcardially with saline and then 4% paraformaldehyde (PFA) in 0.1 M phosphate-buffered saline (PBS). Brains were post-fixed in PFA at 4°C for 48 hours and stored in a cryoprotectant solution. They were then coronally sectioned at 40 μm using a freezing microtome and kept at –20°C until use.

#### 2.5.2. Immunofluorescence

Immunofluorescence assays were performed to study the overexpression of α-syn, its aggregation through phosphorylation (p-syn), the diffusion capacity in the brain of the wild-type and phosphorylated forms of α-syn, and p-syn’s co-expression with dopaminergic neurons (stained with tyrosine hydroxylase, TH). This technique was also used to analyse the astroglial (glial fibrillary acidic protein, GFAP) and microglial (ionised calcium binding adaptor molecule type 1, IBA-1) activation in the dopaminergic nigrostriatal pathway.

All the primary and secondary antibodies used in these stainings, their concentrations and origins are listed in Table S2.

Free-floating sections were washed 3 times (5 minutes each) in 0.1 M PBS. To block non-specific binding sites, sections were then incubated for 2h at room temperature (RT) with agitation in a blocking solution consisting of 5% normal goat serum (NGS) (Sigma-Aldrich, ref. S26-LITER) and 0.5% Triton X-100 in 0.1 M PBS. Sections were then incubated overnight at 4°C with the corresponding primary antibodies (Table S2) in 0.1 M PBS containing 5% NGS and 0.5% Triton X-100. The next day, after 3 5-minute washes in 0.1 M PBS, sections were incubated for 2h with the corresponding fluorescent secondary antibodies (Table S2) in a 0.1 M PBS solution containing 1% NGS and 0.5% Triton X-100. Subsequently, the last 3 5-minute washes were performed in 0.1 M PBS. Finally, the sections were mounted on Superfrost slides (EprediaTM, 12302108) and coverslipped with a mounting medium (Vectashield).

#### 2.5.3. Immunohistochemistry

This technique was applied to evaluate α-syn and p-syn overexpression and dopaminergic neuron integrity via TH staining.

All the primary and secondary antibodies used in these stainings, their concentrations and origins are listed in Table S3.

These immunostains were performed on free-floating sections that were initially washed 3 times (5 minutes each) in 0.1 M PBS. Endogenous peroxidases were quenched with 3% (α-syn and p-syn) or 30% (TH) H_2_O_2_ and 10% methanol in 0.1 M PBS for 30 minutes at RT. After three washes with 0.1 M PBS to block non-specific binding sites, sections were incubated for 1h at RT in a blocking solution containing 5% normal horse serum (NHS) and 0.5% Triton X-100 in 0.1 M PBS. Sections were then incubated overnight at 4°C in the same blocking solution together with the corresponding primary antibody (Table S3). The next day, after 3 5-minute washes in 0.1 M PBS, sections were incubated for 2h at RT with the corresponding biotinylated secondary antibody (Table S3) using 2.5% NHS and 0.5% Triton X-100 in 0.1 M PBS. After three washes in 0.1 M PBS were done, and sections were incubated with an avidin-biotin-peroxidase complex (ABC kit, 1:200, Vector Laboratories, ref. PK-6100) and 0.5% Triton X-100 in 0.1 M PBS for 1h at RT. Peroxidase activity was visualised using the chromogen 3,3′-diaminobenzidine (DAB, Sigma-Aldrich, ref. D5637) and H_2_O_2_. Finally, sections were mounted, air dried for 48h, cleared in xylene for at least 2h and coverslipped with DPX synthetic resin mounting medium (Sigma-Aldrich, ref. 06522).

### 2.6. Quantification procedures

#### 2.6.1. Optical densitometry analysis

Striatal, *nucleus accumbens* (NAcc) and nigral sections of TH; and striatal and nigral α-syn stainings were optically digitised at a resolution of 6400 ppp using an EPSON Perfection V700 scanner and EPSON Scan software (version 3.9.2.2). α-Syn^+^ and TH^+^ fibre densitometry analyses were performed using the ImageJ software (version 2.9.0, NIH). At the 60 days time point, 9-15 striatal, 4-6 NAcc and 4-7 nigral sections of TH per animal were used, and 6-7 striatal, 3-4 nigral sections of α-syn stainings were used. At the 120 days time point, 4-6 striatal, 2-4 NAcc and 5-7 nigral sections of TH per animal were used, and 4-6 striatal, 1-3 nigral sections of α-syn stainings were used. The measured values were normalised for non-specific background staining by subtracting the values obtained from the *corpus callosum* or the basal part of the cerebral peduncle.

#### 2.6.2. Stereological quantification of dopaminergic neurons in the substantia nigra pars compacta

Quantification was performed with an Olympus BX51 microscope and the optical fractionator method via Mercator software (Explora-Nova, La Rochelle, France). This method uses systematic, random sampling of 3D sections to estimate cell numbers. SNc cell counting was done between Bregma –2.70 mm and –3.88 mm, referencing the Paxinos and Franklin atlas (Paxinos and Franklin, 2004).

Cells were counted at 40x magnification within a 4x delineated area, following stereological rules: neurons inside the inclusion lines or touching them were counted, while those on exclusion lines were not. The total number of cells was estimated using the formula: N = ΣQ × (1/ssf) × (1/asf) × (1/hsf), where Q is the number of counted cells. The section sampling factor (ssf) was 1/5, the counting frame size of 50 × 50 μm with a separation between dissectors of 60 μm, and the height sampling fraction (hsf) of 95%. Approximately 5–8 sections per animal were analysed. The volume was estimated using the Cavalieri method to calculate neuron density (cells/mm³) for each group.

#### 2.6.3. Macro-based quantification of axonal swellings

Images were captured using an Olympus BX61 microscope with a 60x oil objective and a z-stack interval of 1 μm, covering the full section thickness. One field of view in the striatum was imaged per hemisphere per coronal section. 10 AAV-α-syn animals (5 males, 5 females) were analysed at each time point, with 6 sections imaged per animal, totaling 12 image stacks per animal.

Axonal swellings in the striatum were quantified using an automated workflow adapted from Quintino et al. (2022). Briefly, a *Batch* of images was generated to undergo the macro-based processing to quantify axonal swellings. The process included converting z-stacks into 8-bit grayscale images, inverting background contrast, and applying several image enhancement steps: rolling ball background subtraction, pixel subtraction, median smoothing, and contrast enhancement. Thresholding was then applied to identify structures.

Only particles larger than 0.53 μm² and with a minimum voxel number of 5 were classified as axonal swellings. For each animal, the total number and median volume of axonal swellings were calculated.

#### 2.6.4. Axonal Degeneration Index calculation

An Axonal Degeneration Index (ADI) was calculated to quantify the extent and progression of axonal damage in the striatum of α-syn mice. This index combined two measures: striatal TH optical density (indicating –a probably linear-fibre loss) and the number of axonal swellings (markers of active degeneration) detected via α-syn immunohistochemistry.

Since axonal swellings follow a non-linear, inverted U-shaped trajectory during degeneration—starting low (or absent) in the early stages, increasing as axons become damaged and eventually declining as fibres fragment and are lost—a continuous index would not accurately reflect degenerative staging.

Instead, an ordinal classification system was used. TH levels and swelling counts were z-score normalised per timepoint, and animals were assigned to one of three degeneration stages: *1)* Early stage (score 1): high TH (z > 0.3), low swelling (z < –0.5); *2)* Intermediate stage (score: moderate TH (z between –0.3 and 0.3), any swelling level; and *3)* Advanced stage (score: low TH (z < –0.3) with either high swelling (z > 0.5, active degeneration) or low swelling (z < 0.5, fibre loss with minimal ongoing degeneration, final stage).

#### 2.6.5. Glial activation

To assess glial activation, GFAP^+^ and IBA-1^+^ cells were analysed in the striatum and SN. Images were acquired with a Zeiss Axioskop fluorescence microscope (40x objective) connected to a Nikon DS-Qi1 mono camera. In each hemisphere, 3 images were taken in the striatum (dorsal, medial, ventral) and 3 in the *substantia nigra* (SN) (medial, central, lateral).

Astrocyte and microglia activation was semi-quantified using ImageJ (version 1.8.0) by calculating the GFAP^+^ and IBA-1^+^ areas relative to the total area (mm²), respectively. To ensure consistent analysis, all images were batch processed using the same settings: background removal with the “Subtract Background” tool (rolling ball radius: 20 pixels), Gaussian Blur (sigma: 2), and automatic brightness/contrast adjustment. The “Moments” thresholding method was applied to select GFAP^+^ and IBA-1^+^ areas. Processed images were saved, and results were exported to a spreadsheet for further analysis.

### 2.7. Statistical analysis

Statistical analyses were conducted in RStudio (version 2024.09.0). Behavioural and morphological data were analysed using a robust two-way ANOVA, implemented via the WRS2 package. The main factors were α-syn group and sex, and an interaction term (α-syn group × sex) was included in the model. Each time point was analysed independently. The results were presented graphically using GraphPad Prism (version 8.0.1; GraphPad Software Inc., USA), with the significance levels represented as follows: “*” denotes the α-syn group effect, with *p<0.05; **p<0.01; ***p<0.001; “#” the sex effect, with #p<0.05; ##p<0.01; ###p<0.001; and “$” denotes the interaction effect (α-syn group × sex), with $p<0.05; $$p<0.01. An independent sample t-test was used for comparisons of the ADI, as only two groups were included in this analysis (A53T α-syn males vs. A53T α-syn females).

## 3. Results

### 3.1. Alpha-synuclein diffusion, overexpression and co-localisation

Stereotaxic injection of AAV A53T human α-syn vectors induced a statistically significant increase in α-syn at the striatal level in both male and female mice. At 60 days post-surgery, two-way robust ANOVA revealed a significant main effect of the α-syn group in both the SN and striatum (Fig. S4A and S4B), indicating increased α-syn levels following AAV-A53T injection. No significant group × sex interaction was observed at this early time point, suggesting similar levels of accumulation in males and females.

At 120 days post-surgery, the main effect of the α-syn group remained significant in both regions (Fig. S4A and S4B). Additionally, a significant group × sex interaction emerged in both the SN and striatum, reflecting a more pronounced accumulation of α-syn in females compared to males at this later stage. This differential progression suggests sex-dependent dynamics in α-syn pathology, with females exhibiting a greater percentage increase in α-syn levels over time.

Next, we assessed whether SNc dopaminergic neurons accumulated pathological α-syn by performing double immunofluorescence for TH and p-syn. Co-localisation of p-syn with TH^+^ neurons confirmed the presence of pathological α-syn within dopaminergic neurons (Fig. S5). P-syn accumulation was restricted to the injection site in the SN and was consistently detected in both sexes across all timepoints (Fig. S6 and S7).

In the ventral tegmental area (VTA), expression of α-syn was observed; however, no p-syn expression was found (Fig. S8A), nor loss of TH^+^ fibers was detected in the NAcc (Fig. S8B), demonstrating the specificity of the model.

### 3.2. Motor impairment in mutant A53T alpha-synuclein mice

In the open field test, total activity was unaffected by A53T α-syn overexpression. However, at 60 days post-injection, α-syn significantly reduced vertical activity and time spent in the central zone (Fig. 1A). A significant α-syn × sex interaction was observed for maximum speed at this time point: A53T α-syn overexpression tended to reduce maximum speed in males while increasing it in females, indicating sex-specific effects on locomotor speed. This interaction was not significant at 120 days post-surgery.

**Fig. 1.**
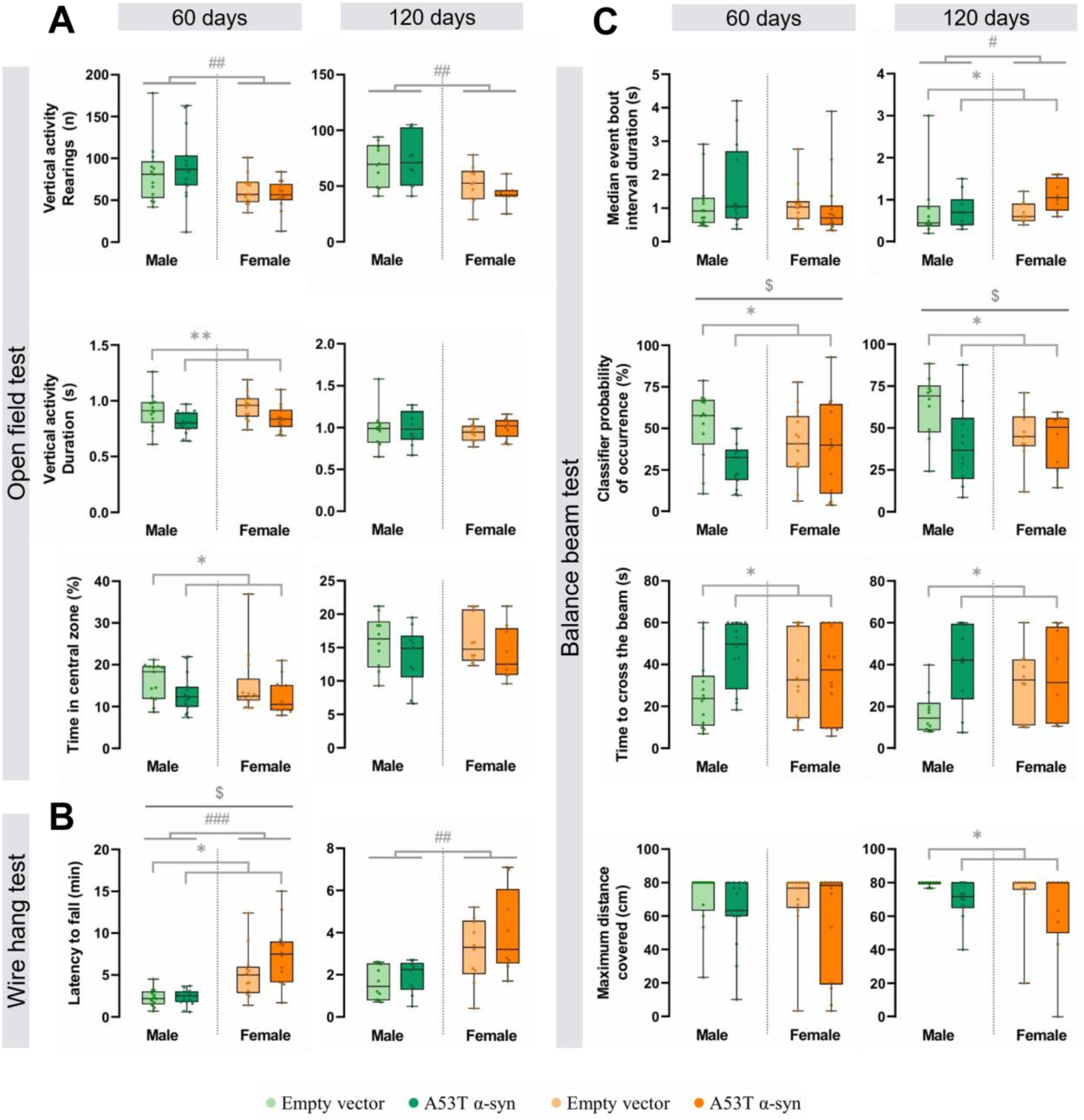
Motor impairment in mutant A53T alpha-synuclein (α-syn) mice. Mice performed the open field **(A)**, wire hang **(B)** and balance beam tests **(C)** at 60 and 120 days post-surgery. Statistical significance was determined using robust two-way ANOVA, with significance levels represented by: * for the α-syn group effect *p<0.05; **p<0.01; # the sex effect #p<0.05; ##p<0.01; ###p<0.001; and $ denoting the interaction effect (α-syn group × sex) with $p<0.05.

Similarly, at 60 days post-surgery, α-syn significantly increased the latency to fall from the grid (p = 0.029), with α-syn animals hanging longer than empty vector mice. This effect was sex-dependent (group × sex, p = 0.047), with α-syn females showing longer latencies (Fig. 1B).

Automated analysis of the balance beam test revealed subtle impairments in motor function and balance in A53T α-syn overexpressing animals at both time points. Specifically, α-syn mice showed reduced walking activity compared to empty vector mice, as indicated by a significantly lower classifier probability of occurrence (p = 0.028 and p = 0.024, respectively) (Fig. 1C). They also failed the test more frequently, receiving the default maximum crossing time (60 s), and took significantly longer to cross the beam (p = 0.039 and p = 0.032, respectively). A near-significant group × sex interaction was noted at both time points for the beam crossing time (p = 0.090 and p = 0.072), suggesting that A53T α-syn overexpression prolonged crossing time primarily in males. At 120 days post-surgery, additional deficits were evident in α-syn mice, including reduced distance travelled along the beam (maximum distance covered, p = 0.046; Fig. 1C) and increased immobility or freezing (median event bout interval duration, p = 0.033; Fig. 1C).

Notably, A53T α-syn overexpression differentially affected sexes across multiple parameters. At both 60 and 120 days post-surgery, α-syn significantly reduced walking percentage in males but not females (classifier probability of occurrence, group × sex, p = 0.047 and p = 0.035, respectively; Fig. 1C). It also increased mean immobility time in males while reducing it in females (mean event bout interval duration, group × sex, p = 0.013). At 120 days, although no differences reached significance, the interaction for immobility duration approached significance (p = 0.066).

No differences were observed at any time point in the pole test (Fig. S9).

Collectively, these results suggest that A53T α-syn overexpression does not affect total activity or gross motor function. However, subtle motor alterations emerge and evolve differently between sexes, ultimately reaching comparable levels of impairment over time.

### 3.3. A53T alpha-synuclein-induced dopaminergic degeneration in the nigrostriatal pathway

To determine whether A53T α-syn overexpression induces dopaminergic degeneration in the nigrostriatal pathway and whether this effect is sex-specific, cell density in the SNc and optical density of TH^+^ fibres in the striatum and *substantia nigra pars reticulata* (SNr) were assessed in both male and female mice.

Stereotaxic injection of AAV-A53T human α-syn did not reduce TH^+^ soma density in the SNc (Fig. 2), nor the TH^+^ dendritic density in the SNr (Fig. S10). However, TH optical density in the striatum decreased following α-syn injection, reaching significance only at 120 days post-surgery, with no significant α-syn × sex interaction (p = 0.294; Fig. 3A).

**Fig. 2.**
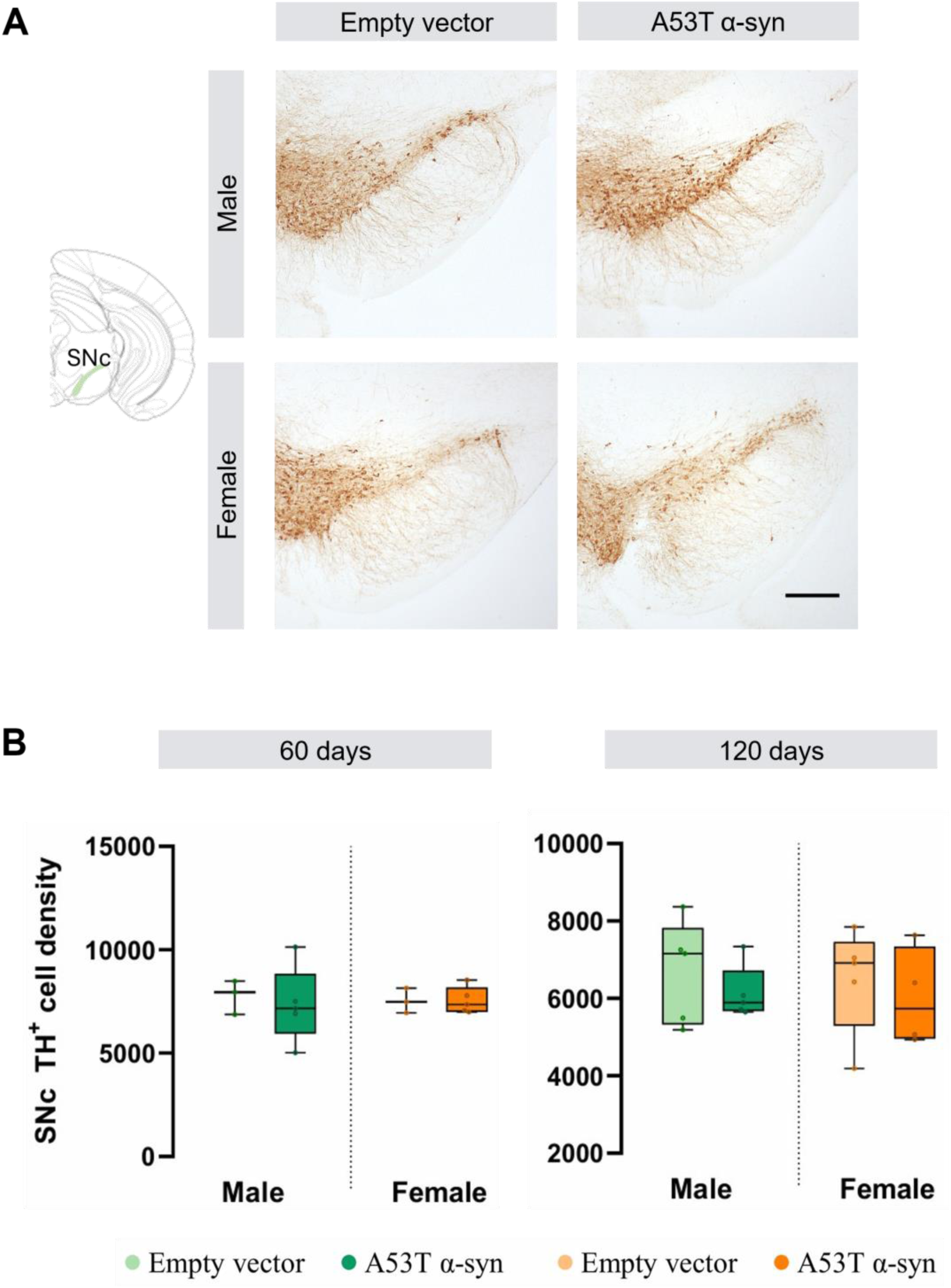
Preserved dopaminergic cell density in the *substantia nigra pars compacta* (SNc) in male and female A53T alpha-synuclein (α-syn) and empty vector animals. (A) Immunohistochemistry for tyrosine hydroxylase (TH) in the *substantia nigra* of representative samples from empty vector and α-syn groups, 60 days after virus administration. Scale bar: 300 μm. **(B)** Stereological quantification of TH^+^ cell density in the SNc at 60 and 120 days post-surgery.

**Fig. 3.**
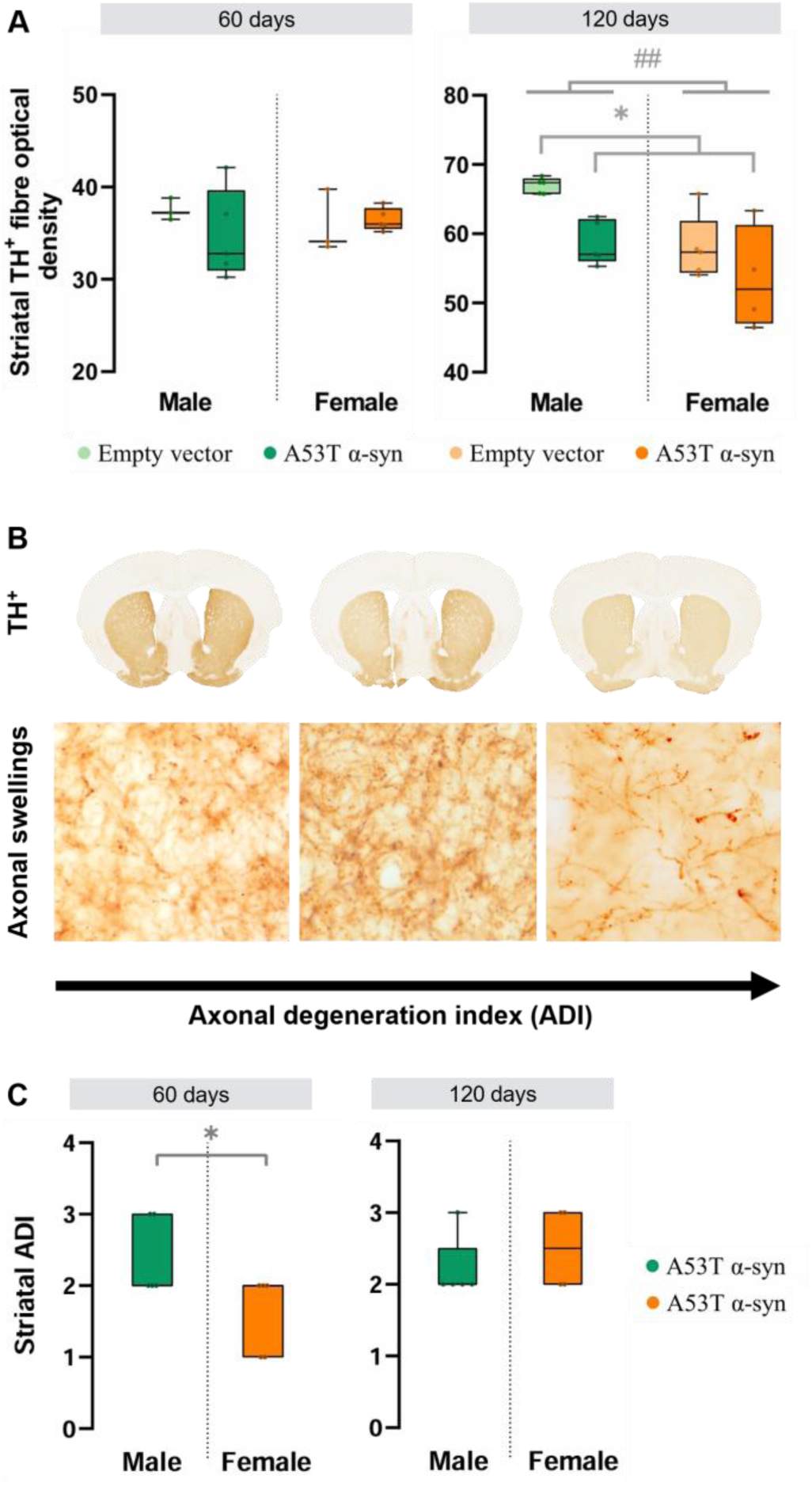
Sex-dependent pattern of striatal degeneration in A53T alpha-synuclein (α-syn) mice. **(A)** Quantification of the optical density of dopaminergic fibres in the striatum in male and female A53T α-syn and empty vector animals. Dopaminergic neurons were positive for tyrosine hydroxylase (TH^+^). Statistical significance was determined using robust two-way ANOVA, with significance levels represented by: * for the α-syn group effect *p<0.05; and # the sex effect ##p<0.01. **(B)** The Axonal Degeneration Index (ADI) was calculated to quantify the extent and progression of axonal damage in the striatum of α-syn mice. This index was derived by combining two complementary measures: striatal TH optical density and the number of striatal axonal swellings detected via immunohistochemistry against α-syn. **(C)** Striatal ADI in male and female A53T α-syn mice. Statistical significance was determined using an independent sample t-test, with *p<0.05.

Importantly, TH optical density alone may underestimate early axonopathy, as TH^+^ fibres can exhibit swelling and fragmentation prior to complete loss of TH expression. These pathological changes often peak before fibre loss occurs, leading to potential underestimation of degeneration in the presence of prominent axonal swellings. Therefore, A53T α-syn mice were classified into three stages based on combined measures of TH optical density and axonal swelling counts in the striatum (Fig. 3B). When accounting for axonal swelling, α-syn males exhibited significantly more advanced TH axonal degeneration than females at 60 days post-surgery (p = 0.020, Fig. 3C), revealing sex differences in the neurodegenerative progression despite converging to similar levels of striatal degeneration by 120 days post-surgery.

### 3.4. Neuroinflammation in the nigrostriatal pathway

Next, we investigated whether overexpression of α-syn induced glial activation of both astrocytes (labelled with GFAP) and microglia (labelled with IBA-1) in the striatum and SN of A53T α-syn mice. At 60 days post-surgery, α-syn animals showed significantly increased astrocytic activation in the striatum compared to empty vector controls (p = 0.009; Fig. 4). This elevated astrocytic response persisted at 120 days (p = 0.029; Fig. 4), indicating sustained neuroinflammation. Representative images of astrocytic activation in the striatum and SN are shown in Fig. 4. In contrast, AAV-A53T α-syn injection into the SNc did not induce significant astrocytic activation in the SN at either time point (Fig. 4), nor did it trigger microglial activation in the striatum or SN at any time. No sex-specific differences were observed in either marker.

**Fig. 4.**
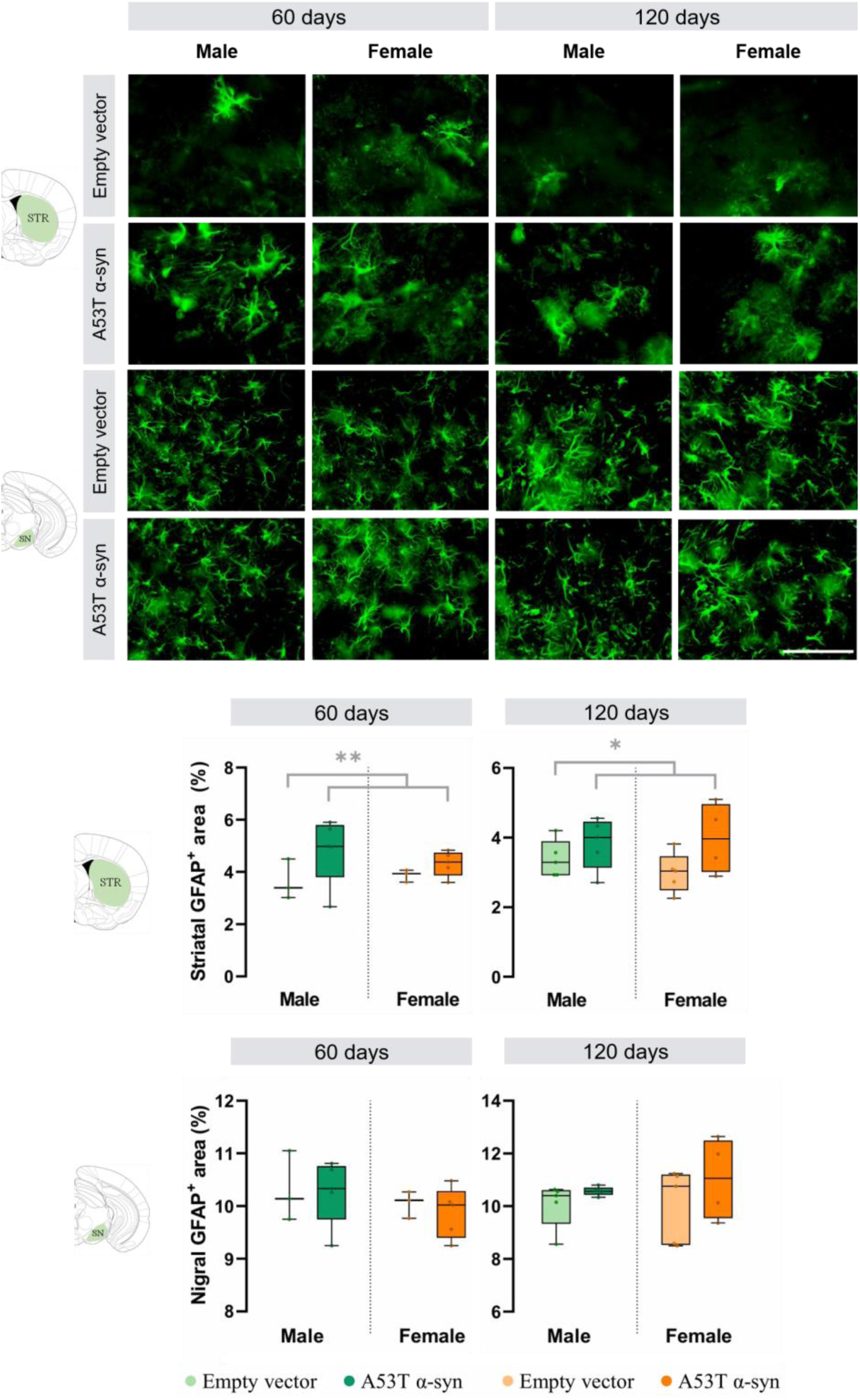
Neuroinflammation in the nigrostriatal pathway. Representative images of striatal and nigral astrocytic activation in empty vector and alpha-synuclein (α-syn) animals at 60 and 120 days after surgery. Scale bar: 80 µm. Quantification of glial fibrillary acidic protein (GFAP) expression in the striatum and substantia nigra at both time points. Statistical significance was determined using robust two-way ANOVA, with significance levels represented by: * for the α-syn group effect *p<0.05; **p<0.01.

## 4. Discussion

In this study, we examined the consequences of mutant A53T α-syn overexpression in a mouse model, focusing on the temporal progression of pathological changes and their sex-specific manifestations. The key pathological domains assessed included motor function, dopaminergic fibre integrity, axonal pathology, and astrocytic activation. Our findings demonstrate that while both sexes eventually reach comparable levels of neurodegeneration, the timing and trajectory of these processes differ significantly between males and females, particularly at early stages.

Behaviourally, A53T α-syn overexpression induced early motor and affective disturbances detectable at 60 days post-injection. Mice exhibited increased anxiety-like behaviour in the open field test, consistent with clinical observations in PD where anxiety frequently coexists with motor deficits and may precede overt motor symptoms (Coakeley et al., 2014). Despite preserved gross motor function in the pole test, subtle impairments were detected in motor coordination as assessed by the balance beam test, reinforcing the notion that balance and gait disturbances may emerge prior to more global motor deficits (Costa et al., 2020; Sirabella et al., 2018). Notably, sex-specific differences emerged across multiple behavioural domains. In the wire hang test, females exhibited significantly longer latency to fall than males at both 60 and 120 days, which may reflect pathological hypertonia—an early manifestation of PD—leading to sustained muscular rigidity and prolonged grip. Alternatively, this behaviour may also be modulated by heightened affective responses, as increased anxiety may intensify avoidance of falling, despite no overt sex differences in open field anxiety measures. Importantly, our automated analysis of balance beam walking behaviour, utilising DeepLabCut and SimBA (Bidgood et al., 2024), revealed subtle but consistent motor deficits in males, including longer immobility and reduced active walking time compared to females. This supports the presence of early, sex-dependent motor dysfunction that might be overlooked by traditional manual scoring. The literature on sex differences in early motor phenotypes in PD remains inconclusive. Some clinical studies suggest milder motor symptoms in men (Solla et al., 2012), while others report less severe impairment in women (Kang et al., 2022; Kim et al., 2018). The reduced vulnerability in females may stem from higher physiological dopamine levels in the nigrostriatal pathway (Lee et al., 2015) and the neuroprotective actions of estrogens via anti-inflammatory, antioxidant, anti-apoptotic, and anti-aggregation mechanisms (Rajsombath et al., 2019; Rugbjerg et al., 2013; Thadathil et al., 2021). However, sex-specific data on early-stage PD remain limited, underscoring the novelty of our findings.

Morphological analyses further corroborated the temporally distinct progression of neurodegeneration between sexes. While striatal TH optical density exhibited a modest decline at 60 days, significant reduction was only observed by 120 days. However, early axonal swelling and fragmentation may confound isolated TH optical density measurements by temporarily maintaining TH positivity despite structural disruption. By incorporating both TH optical density and axonal swelling counts into a composite axonal degeneration index (based on z-scoring both variables and stratifying animals into early, intermediate, and advanced degeneration stages), we revealed that A53T α-syn male mice exhibited more pronounced neurodegenerative changes as early as 60 days post-injection. Importantly, both males and females exhibited comparable TH levels in the SNr and similar dopaminergic cell densities in SNc to empty vector mice at these time points, suggesting that axonal pathology precedes measurable loss of dopaminergic terminals or cell bodies and highlights sex-specific differences in the early pattern of striatal neurodegeneration that are not detectable by TH optical density measures alone. Consistent with our observations, previous work in A53T α-syn mice demonstrated dopaminergic neuron loss in the SNc only at later stages (12 months), with males exhibiting earlier gross motor impairment (Costa et al., 2020; Sirabella et al., 2018). *Postmortem* analyses in PD patients similarly indicate that at the onset of motor symptoms, the extent of dopaminergic terminal loss in the striatum or putamen exceeds neuronal loss in the SNc (Bernheimer et al., 1973; Kish et al., 1988; Scherman et al., 1989). While approximately 60–80% of SNc dopamine neurons are lost by late disease stages (Fearnley and Lees, 1991; Pakkenberg et al., 1991), the depletion of dopaminergic markers in the striatum is even more severe, emphasizing that axonal degeneration precedes somatic neuronal loss (Burke and O’Malley, 2013). Human neuroimaging studies have further demonstrated that dopaminergic neurodegeneration initially manifests more prominently in axonal projections (Caminiti et al., 2017), which is consistent with our findings. Our data thus provide preclinical support for this axonopathy-first model, while introducing novel evidence of sex-dependent variations in this retrograde degenerative process.

Astrocytic activation was evident in the striatum at both 60 and 120 days, aligning temporally with dopaminergic axonal pathology. No significant astrocytic changes were detected in the SN, which may reflect a later emergence of glial responses in regions spared from early dopaminergic loss. Previous reports of SN astrogliosis in 12-month-old A53T mice (Costa et al., 2020; Sirabella et al., 2018) support this notion. Interestingly, while no sex differences in astrocytic activation were detected at our time points, it remains plausible that earlier assessments might uncover divergent glial responses between males and females. Microglial activation, which often precedes astrogliosis, was not detected at these time points, but may have occurred transiently at earlier stages, initiating sustained astrocytic activation in response to accumulating α-syn pathology. The potential existence of temporally and sexually divergent glial responses warrants further investigation.

This study has several strengths. First, by integrating both TH optical density and axonal swelling counts into a composite degeneration index, we were able to sensitively capture early neurodegenerative changes that might be missed by traditional single-parameter assessments. Second, the application of machine learning-based automated behavioural analysis (DeepLabCut and SimBA) allowed for high-resolution detection of subtle motor impairments, minimising observer bias and enhancing reproducibility. These methodological innovations provide a more nuanced characterisation of early disease stages, critical for understanding sex-specific trajectories in PD progression.

Nevertheless, some limitations should be acknowledged. Due to technical constraints, only two time points (60 and 120 days post-injection) were assessed, precluding a finer resolution of early dynamics. The inclusion of earlier time points could help clarify the initiation of glial responses and better delineate sex-specific temporal differences. Similarly, assessments at later stages may reveal whether dopaminergic neuronal loss in the SNc diverges further between sexes. Moreover, while AAV-mediated models provide valuable insights, they inherently exhibit variability in viral transduction efficiency; although animals with insufficient viral spread were excluded, this remains a general limitation of the model. Finally, a more comprehensive statistical design incorporating time, sex, and treatment factors would enhance future studies.

## 5. Conclusion

In summary, our findings demonstrate that A53T α-synuclein overexpression induces early, progressive dopaminergic axonopathy associated with subtle but significant sex-specific differences in motor and neuroinflammatory profiles. Males exhibited earlier and more severe axonal pathology, while females demonstrated possible hypertonia-associated motor behaviours. These results emphasize the importance of considering sex as a biological variable in PD models and highlight the need for sex-specific therapeutic approaches targeting early disease stages.

## Acknowledgements

We would like to thank Dr. Benjamin Dehay (Univ. Bordeaux, CNRS, IMN, UMR 5293, F-33000, Bordeaux, France) for providing the viruses used in this work, as well as Dr Nicolas Mallet (Univ. Bordeaux, CNRS, IMN, UMR 5293, F-33000, Bordeaux, France) for helpful discussion.

## Author contributions

T.M.H. designed the study and the experimental strategy; A.M.G., J-V.L. and T.M.H. provided oversight and guidance; M.Z. and J-A.R.O. performed the surgeries; M.Z. and R.B. performed the experiments and prepared the figures; A.M.G. carried out the statistical analyses and wrote the original draft; M.Z., R.B., A.M.G. and T.M.H. edited the manuscript. All the contributors revised critically and gave their approval to the final version of the manuscript.

