## Supplementary information for "Sex-specific vulnerability to behavioural and morphological alterations in a mouse model of Parkinson’s disease overexpressing mutant A53T alpha-synuclein"

**<u>CONTENTS:</u>**

**Supplementary Material and Methods**

**Supplementary Tables (3)**

**Supplementary Figures (10)**

**Supplementary Information References**

* Corresponding author:

**Raphaelle Bidgood**
Department of Pharmacology, Faculty of Medicine and Nursing, University of the Basque Country (EHU)
Postal address: Department of Pharmacology, Faculty of Medicine and Nursing, University of the Basque Country (EHU), Barrio Sarriena s/n, 48940-Leioa, Spain
Telephone number: +33 695695364
ORCID number: 0009-0005-8190-1461

### Experimental design

**Table S1. Summary of the sample sizes used in the behavioural and morphological studies.** Both studies were conducted at 60 and 120 days post-surgery on male and female mice that had been injected with adeno-associated viral (AAV) vectors encoding either A53T mutant human alpha-synuclein (α-syn) or the empty vector.

| | Behavioural study | Morphological study |
|---|---|---|
| **60 days** | Male α-syn (n = 15)<br>Male empty vector (n = 13) | Male α-syn (n = 5)<br>Male empty vector (n = 3) |
| | Female α-syn (n = 14)<br>Female empty vector (n = 13) | Female α-syn (n = 5)<br>Female empty vector (n = 3) |
| **120 days** | Male α-syn (n = 10)<br>Male empty vector (n = 10) | Male α-syn (n = 5)<br>Male empty vector (n = 5) |
| | Female α-syn (n = 9)<br>Female empty vector (n = 10) | Female α-syn (n = 4)<br>Female empty vector (n = 5) |

### Behavioural assessment

The behavioural assessment carried out included the open field test, wire hang test, balance beam test and pole test (Fig. S1).

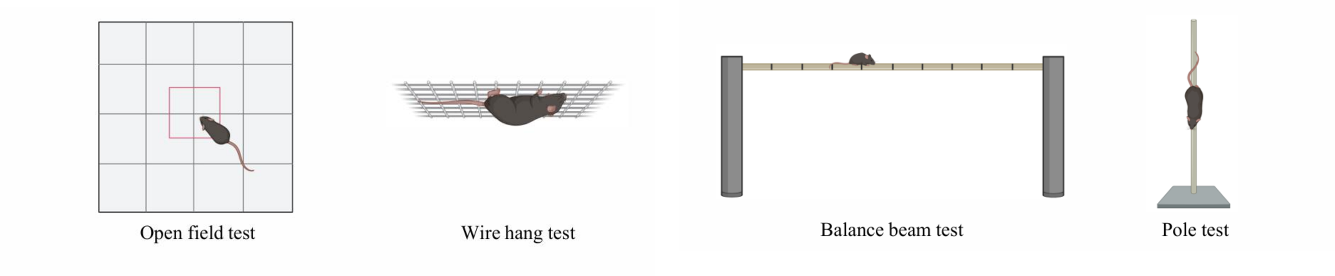


**Fig. S1. Illustrative images of the behavioural tests conducted in this study.** The open field, wire hang, balance beam and pole tests were all performed at both 60 and 120 days post-surgery.

#### *Open field test*

The open field test is the simplest locomotor activity test, which assesses general spontaneous motor activity. A mouse is placed in the centre of an open arena and its movements are recorded, including the number and duration of rearings, resting time and total activity (Brooks and Dunnett, 2009). Activity detection was fully automated using Actitrack software (Panlab, Spain), which was specially developed for this purpose. This apparatus consists of a square arena (44 x 44 x 35 cm) with parallel infrared beams spaced 2.5 cm apart to accurately detect animal movement. Mice were placed in the centre of the arena and were allowed to explore the arena for 10 minutes (period of acclimatisation) before measurements were taken for a further 10 minutes. After each session, the arena was cleaned with 70° alcohol and the sawdust was replaced before another animal was placed into the arena. The computer control unit analysed various parameters, such as the total activity, total distance travelled, resting time, vertical activity, speed and time spent performing fast (more than 5 cm/s), slow (less than 5 cm/s) and stereotyped movements. The software also enabled the arena floor to be divided into two squares to distinguish between the central and peripheral areas. This division made it possible to quantify the relative distance in the central area using the following formula: 100 x [total distance covered in the centre / total distance covered in the periphery].

#### *Wire hang test*

The wire hang test (grid test) is used to assess motor coordination and muscle strength. Mice were placed onto a standard cage grid, which was gently shaken to enable the animals to grasp the wires, before turning the grid upside down. The grid was positioned approximately 50 cm above a standard sawdust-filled cage. This test was performed on two consecutive days, with the latency time for each animal to fall into the cage bedding measured each day. The maximum time allowed for each test was 15 minutes.

***Balance beam test***

The beam-walking test in rodents offers the possibility of measuring subtle deficits in motor skills and balance that other motor tests, such as the rotarod, may not be able to detect. The task assesses an animal's ability to walk across a wooden beam 80 cm long and 1 cm wide at a height of 60 cm (Fig. S2A). The mouse was placed in front of the start line and was removed from the beam either when it had crossed the finish line or when the time on the beam exceeded 1 minute. Animals were given 3 trials, each separated by 10 minutes, and the mean of the performance of each of the 3 trials was calculated for each parameter studied. Prior to the first trial, mice were acclimatised to the behavioural room for 30 minutes. One day prior to testing, mice were trained to cross the beam under the same conditions as the test (3 trials per animal, with a 10-minute interval between each trial). Only test sessions were recorded for in-depth analyses combining manual and automated methods.

Each trial was recorded by two cameras for manual and automated analysis (cameras 1 and 2, respectively). Camera 1 was placed at the rear of the beam to provide a back view of the mouse crossing the beam for the manual quantification of hind paw slips. A slip was defined as the foot leaving the top of the beam (Fig. S2B). Camera 2 was placed laterally to the beam to capture a side view of the full length of the beam, for automated measurement of the time to cross the beam, the maximum distance travelled (using region of interests, ROIs) and an in-depth analysis of mouse walking behaviour. Videos recorded by camera 2 were pre-processed to show only the entirety of the trial (beginning when the animal crossed the start line).

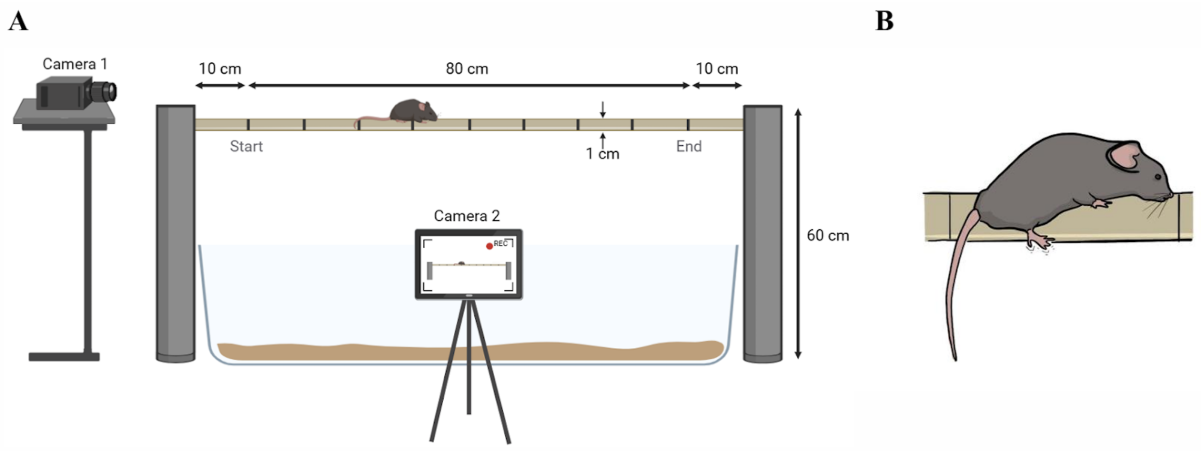


**Fig. S2. Illustrative images of the balance beam test. (A)** Schematic representation of the experimental setup for the balance beam test, which assesses the mouse's ability to walk across a wooden beam (80 cm long and 1 cm wide at a height of 60 cm). Cameras 1 and 2 (placed at the rear and laterally to the beam, respectively) recorded each trial for in-depth manual and automated analyses, post-behavioural sessions. **(B)** Illustration of a hind paw slip, defined as the back foot leaving the top of the beam (Bidgood et al., 2024).

The automated analysis followed the pipeline previously published by Bidgood et al. (2024), using the open-source tools DeepLabCut (version 2.2.3; Nath et al., 2019) and SimBA (version 2.8.5; Goodwin et al., 2024). Briefly, DeepLabCut enabled the automated tracking of the animal along the beam using markerless pose estimation (with 5 keypoints: "Nose", "Head", "Bodytop", "Bodymiddle" and "Tailbase"), while SimBA facilitated the automated behavioural classification to detect mouse walking patterns. The SimBA ROI interface was used to draw the 10 ROIs along the beam, each one delimited by hand-drawn lines marking every 10 cm on the beam. SimBA provided the number of entries in each ROI as well as the time spent in each one (considering the "Bodymiddle" body-part as the reference). The ROI analysis was used to determine the time taken to cross the beam (s) and the maximum distance covered (cm) in an automated manner. Animals entering a ROI at least once were considered to have crossed the corresponding line, aided to calculate the maximum distance covered (i.e. the distance associated with the furthest line from the start line that was crossed by the animal). Values of the times spent in each ROI of the beam were summed to determine the time to cross the beam.

Videos of mice that never crossed the start line were excluded from both the manual (slip counts) and automated (with DeepLabCut and SimBA) analyses, as these subjects never walked along the beam. For the ROI analysis, videos excluded from the DeepLabCut analysis were assigned the default values of a failed trial (60 s to cross the beam and 0 cm as the maximum distance covered), as these animals never crossed the start line.

***Pole test***

The pole test is used to assess motor dysfunction. Mice were placed facing upwards to the top of a 55 cm long, 1 cm diameter pole (Fig. S3). The time taken to make a 180° turn (latency to turn downwards), the descent time and total time spent on the pole were measured. The test was performed on two consecutive days: three trials were performed on the first day (training), followed by three trials on the second day (final). The final trials were recorded with a front-view camera for subsequent manual analysis.

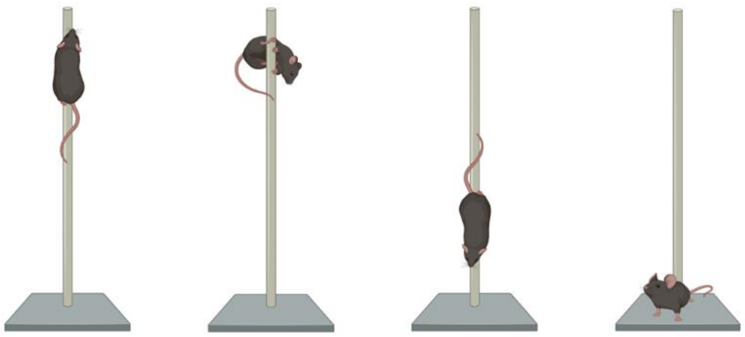

**Fig. S3. Illustrative images of the pole test.** The mouse is first placed on the top of the pole, facing upwards, marking the start of the test. Using a manual chronometer, the time the mouse takes to turn downwards is measured, as well as the subsequent descent time and the total time spent on the pole. The test ends once the animal reaches the bottom of the pole. Adapted from BioRender.

### Immunoassays

#### *Immunofluorescence*

**Table S2. Primary and secondary antibodies used in immunofluorescence assays, concentrations used and origin.**

| | | |
|---|---|---|
| **Primary antibodies** | | |
| Mouse anti-α-syn | 1:10000 | Abcam (Ref. AB27766) |
| Rabbit anti-p-syn | 1:1000 | Abcam (Ref. AB51253) |
| Mouse anti-TH | 1:1000 | Merck Millipore (Ref. MAB5280) |
| Rabbit anti-IBA-1 | 1:1000 | Fujifilm Wako (Ref. 019-19741) |
| Rabbit anti-GFAP | 1:500 | Dako (Ref. Z0334) |
| **Fluorescence secondary antibodies** | | |
| Goat anti-mouse Alexa Fluor 594 | 1:400 | Invitrogen (Ref. A11005) |
| Goat anti-rabbit Alexa Fluor 594 | 1:400 | Invitrogen (Ref. A11012) |
| Goat anti-rabbit Alexa Fluor 488 | 1:400 | Invitrogen (Ref. A11008) |
| Goat anti-mouse Alexa Fluor 488 | 1:400 | Invitrogen (Ref. A11001) |

#### *Immunohistochemistry*

**Table S3. Primary and secondary antibodies used in immunohistochemical assays, concentrations used and origin.**

| | | |
|---|---|---|
| **Primary antibodies** | | |
| Mouse anti-α-syn | 1:10000 | Abcam (Ref. AB27766) |
| Rabbit anti-p-syn | 1:1000 | Abcam (Ref. AB51253) |
| Rabbit anti-TH | 1:1000 | Merck Millipore (Ref. AB152) |
| **Biotinylated secondary antibodies** | | |
| Horse anti-mouse IgG | 1:200 | Vector Laboratories (Ref. BP-2000-50) |
| Horse anti-rabbit IgG | 1:200 | Vector Laboratories (Ref. BA1100-1.5) |

**Alpha-synuclein diffusion, overexpression and co-localisation**

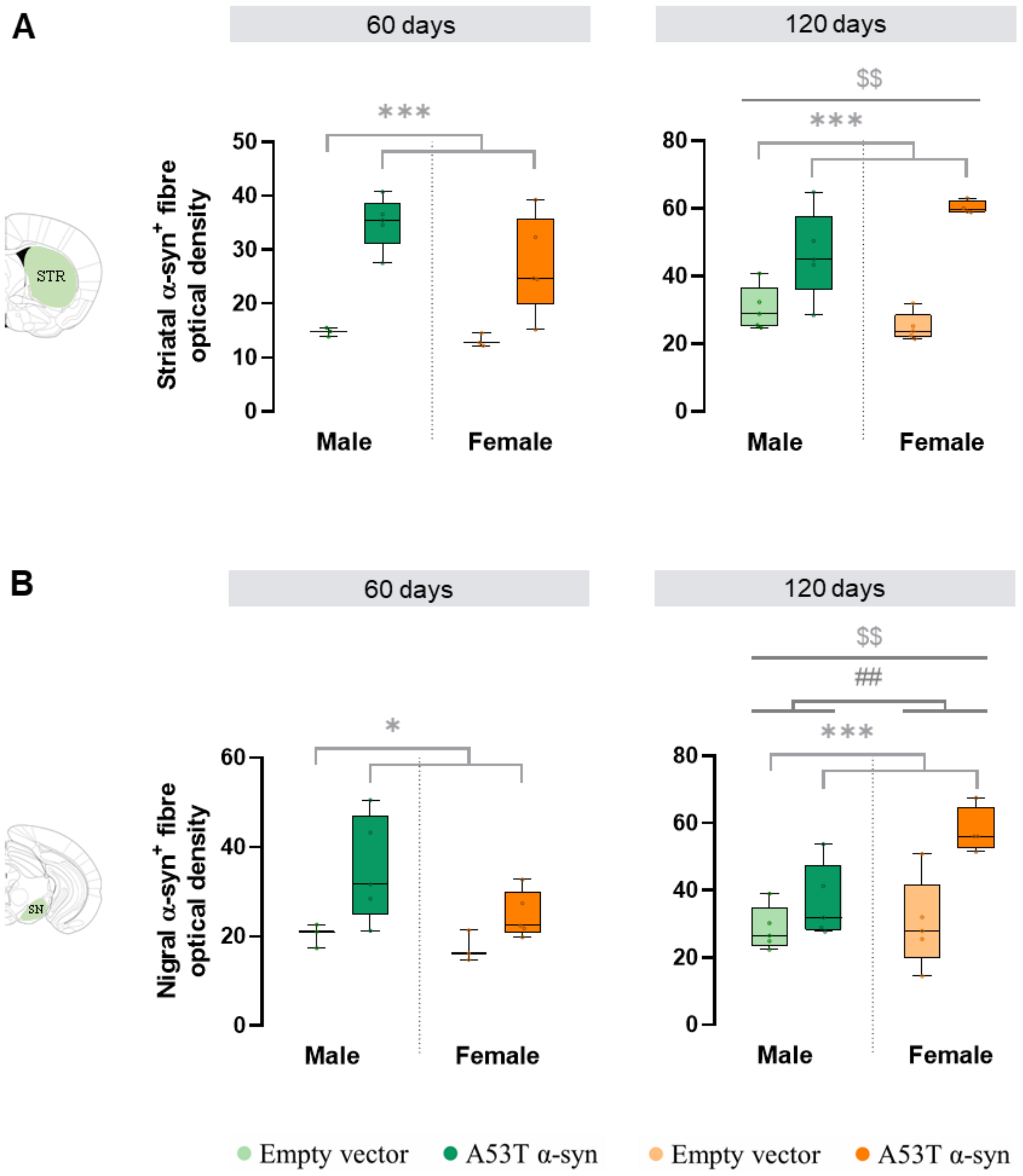


**Fig. S4. Alpha-synuclein (α-syn) overexpression in male and female A53T α-syn and empty vector animals.** Quantification of the optical density of α-syn$^+$ fibres in the striatum **(A)** and *substantia nigra* (SN) **(B)** in male and female mice at both experimental time points: 60 and 120 days post-injection. Statistical significance was determined using robust two-way ANOVA, with significance levels represented by: * for the α-syn group effect *p<0.05; ***p<0.001; # the sex effect ##p<0.01; and $ denoting the interaction effect (α-syn group × sex) with $$p<0.01.

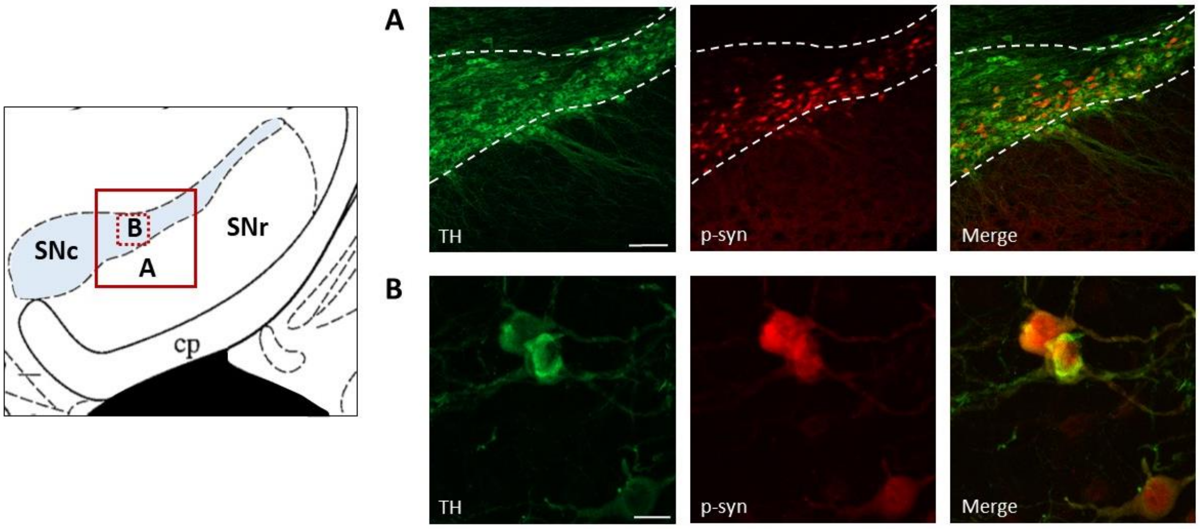


**Fig. S5. Representative images of the *substantia nigra pars compacta* (SNc) showing the co-expression of phosphorylated alpha-synuclein (p-syn) and tyrosine hydroxylase (TH).** Positive labelling is shown for TH (in green), p-syn (in red) and their co-expression (in yellow) in dopaminergic neurons of the SNc. **(A)** Scale bar: 100 µm (objective 10x). **(B)** Scale bar: 15 µm (objective 63x).

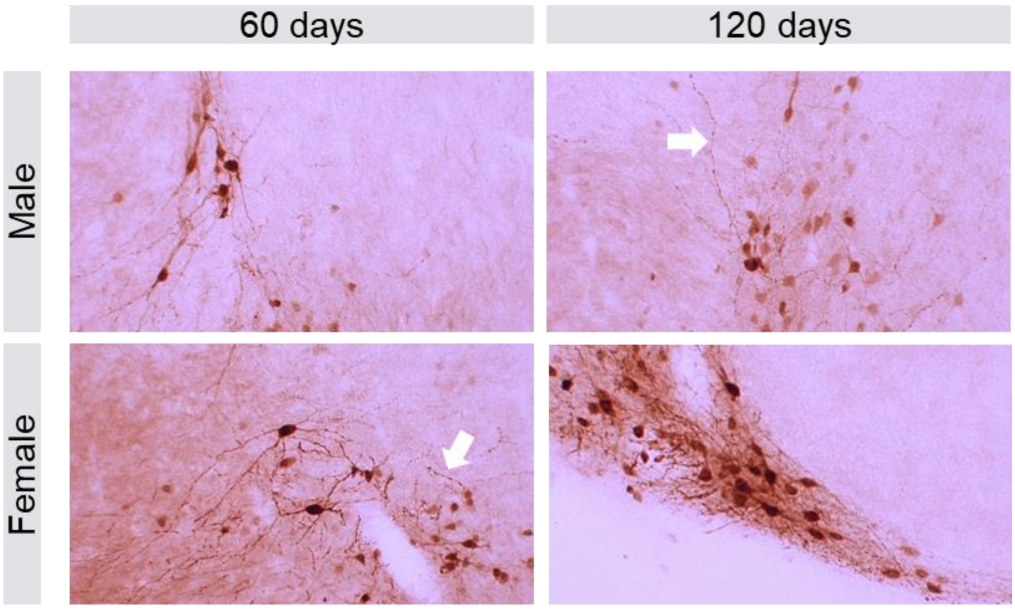


**Fig S6. Expression of phosphorylated alpha-synuclein (p-syn) in the *substantia nigra* of male (top) and female (bottom) mice at 60 and 120 days after surgery.** White arrows indicate the axonal swellings observed in these p-syn-expressing neurons.

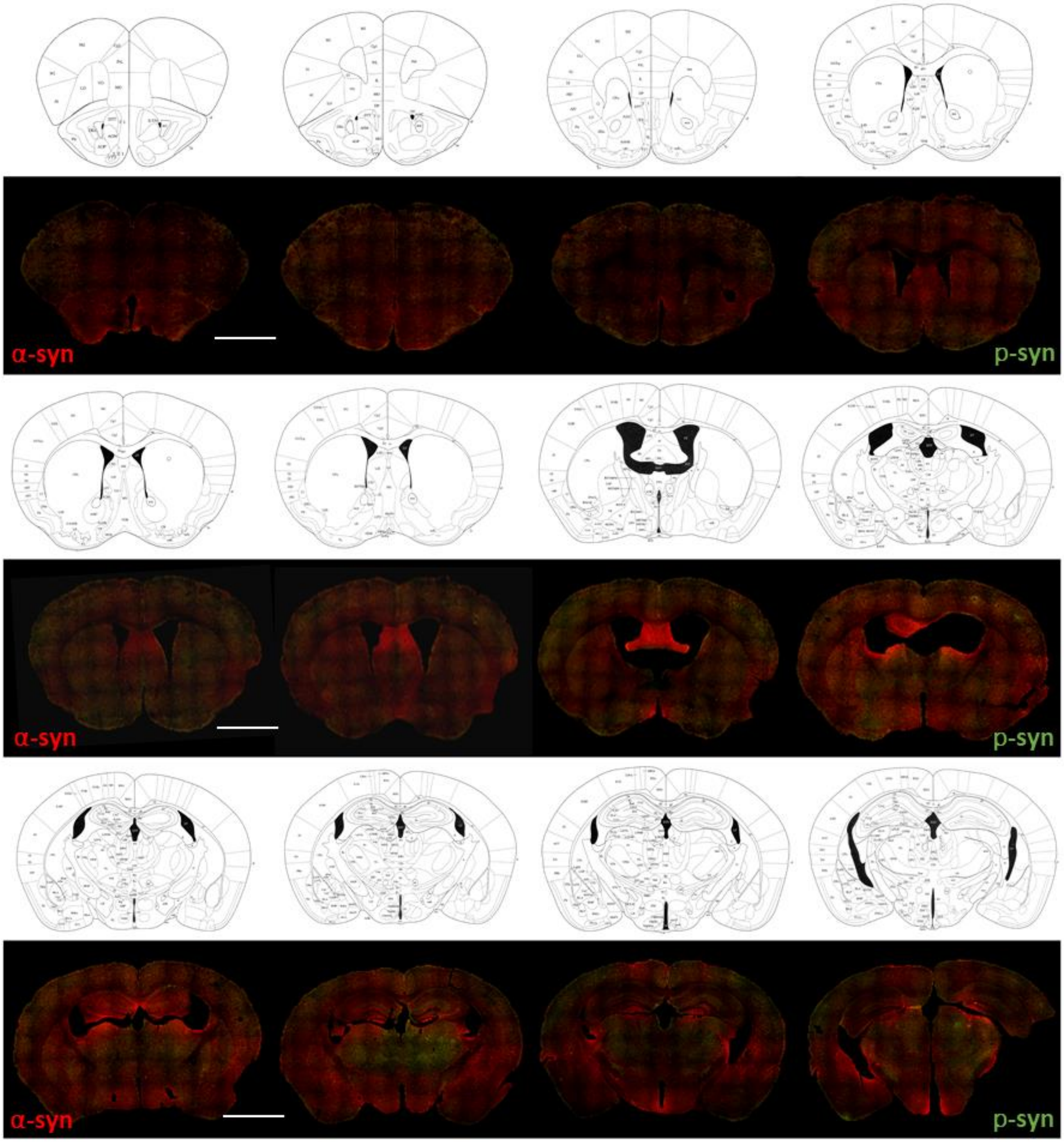
α-syn
p-syn
α-syn
p-syn
α-syn
p-syn

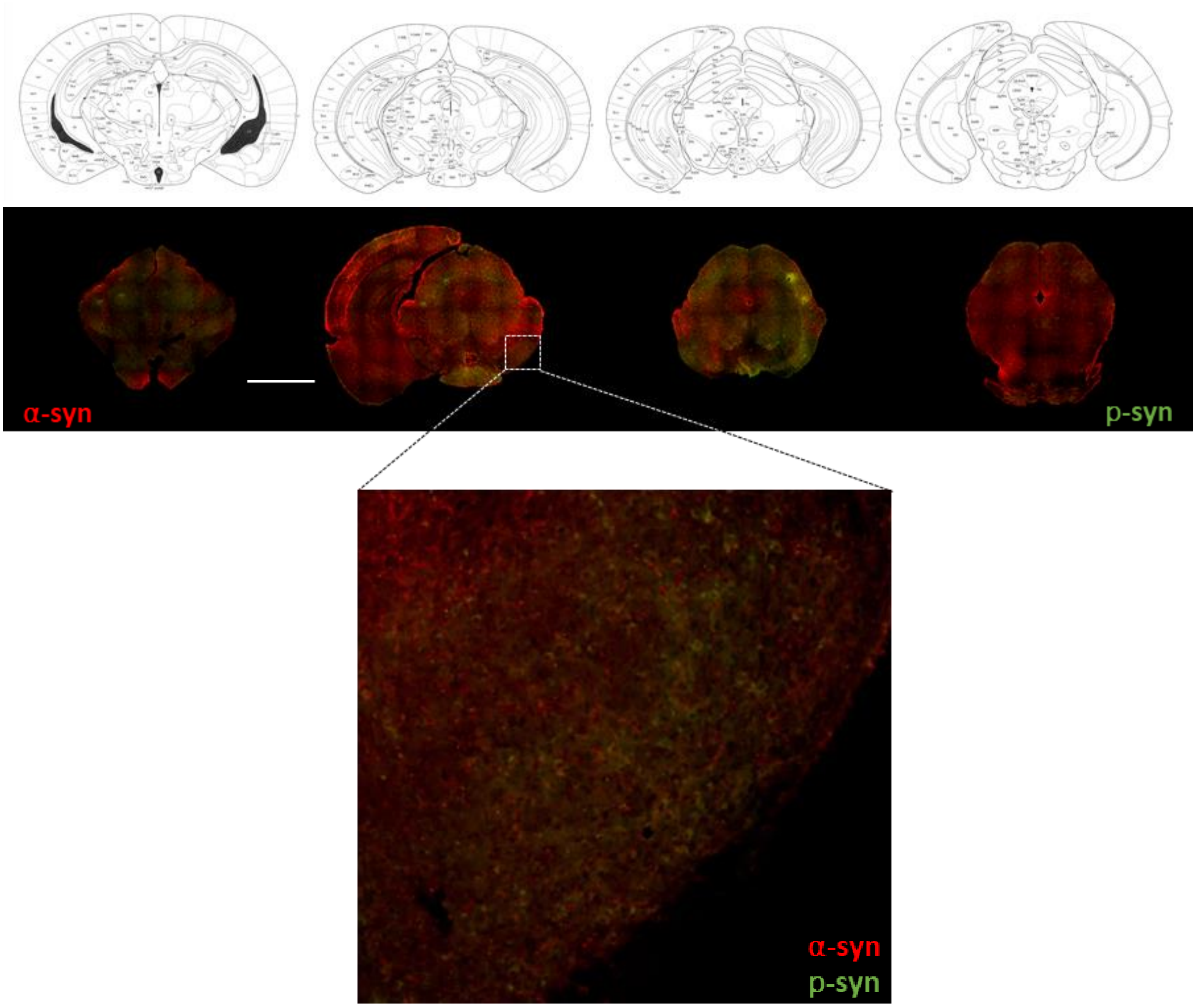


**Fig. S7. Serial immunofluorescence for alpha-synuclein (α-syn, in red) and phosphorylated alpha-synuclein (p-syn, in green).** Serial brain sections from a male animal 120 days after surgery. Scale bar: 2000 μm.

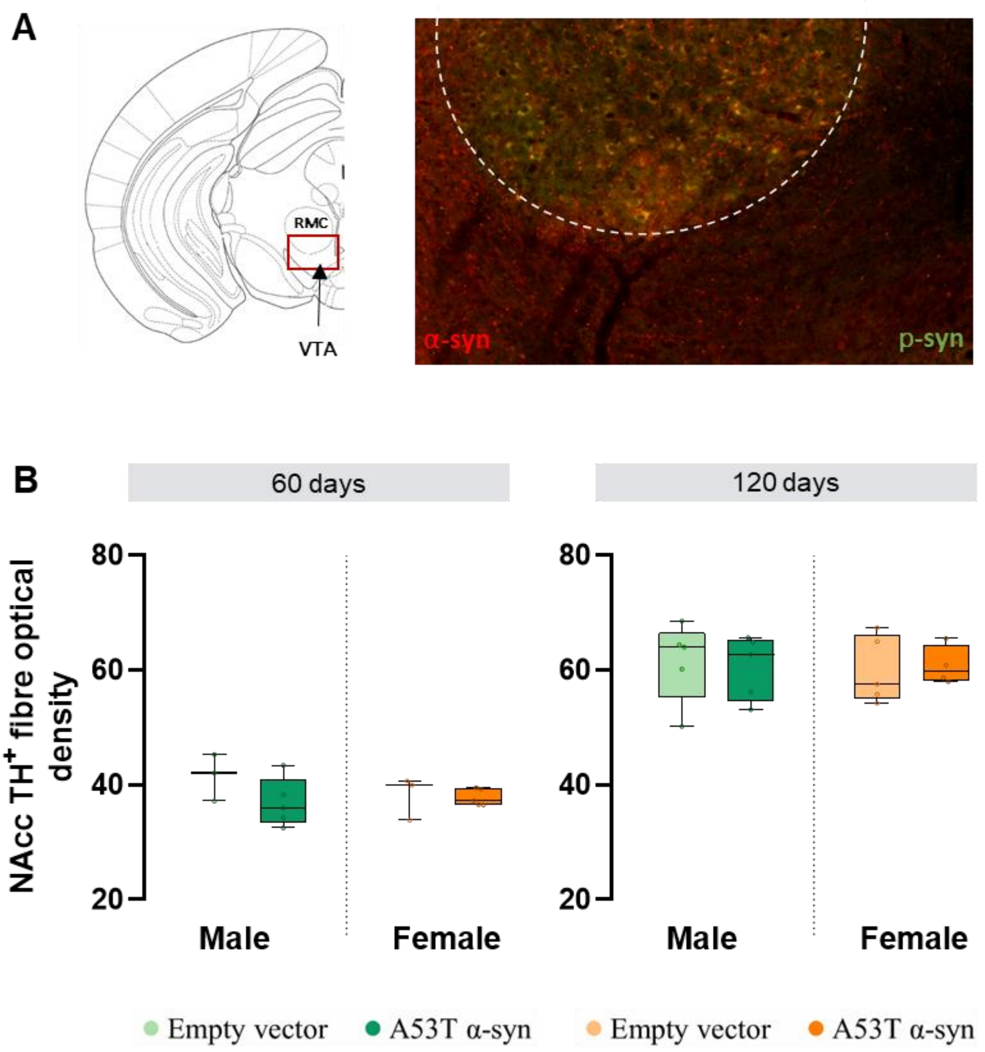


**Fig. S8. Preserved dopaminergic projections from the ventral tegmental area (VTA) to the *nucleus accumbens* (NAc). (A)** Double immunofluorescence for alpha-synuclein (α-syn, in red) and phosphorylated alpha-synuclein (p-syn, in green) of a male animal 120 days after surgery. **(B)** Quantification of the optical density of dopaminergic fibres in the NAcc in male and female A53T α-syn and empty vector animals. Dopaminergic neurons were positive for tyrosine hydroxylase ($TH^+$). No differences in dopaminergic degeneration were observed in the NAcc at either 60 days or 120 days post-surgery.

**Motor impairment in mutant A53T alpha-synuclein mice**

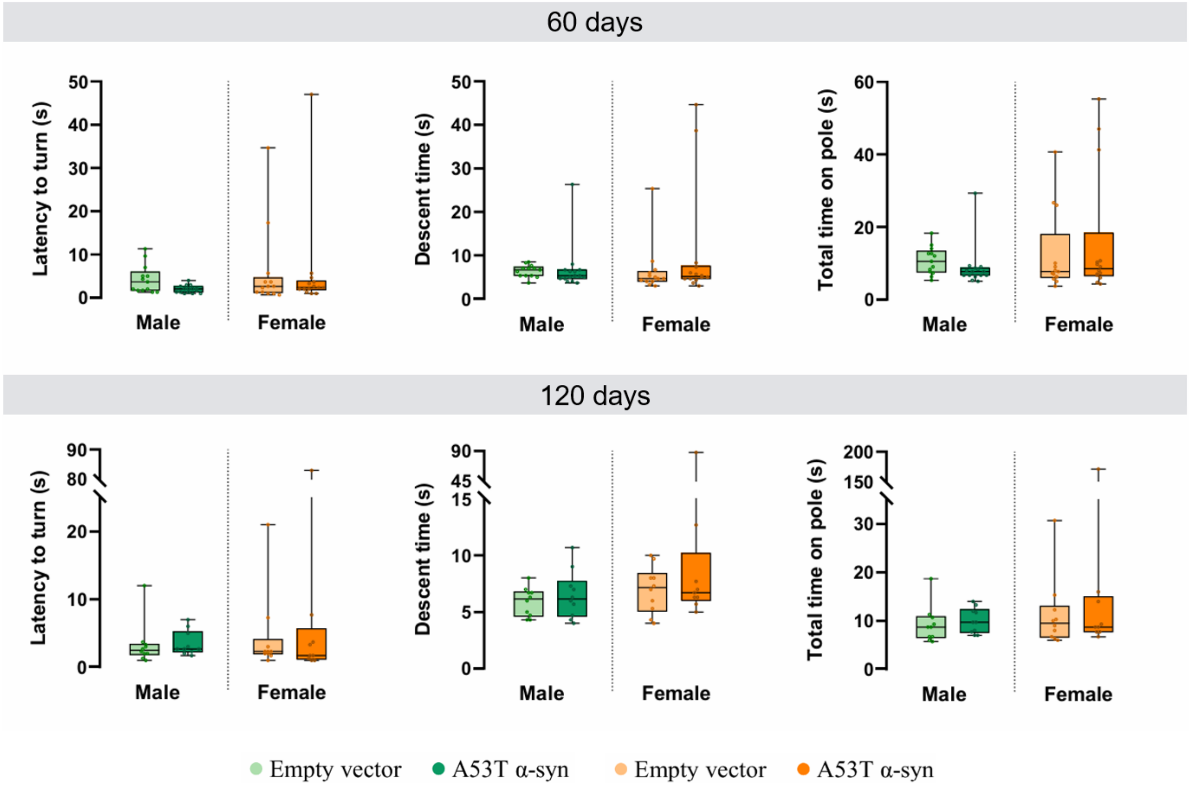


**Fig. S9. Pole test in male and female A53T alpha-synuclein (α-syn) and empty vector animals.** Mice performed the test at 60 and 120 days post-surgery. No statistically significant differences were found in the latency to turn, the descent time, and total time spent on the pole.

**A53T alpha-synuclein-induced dopaminergic degeneration in the nigrostriatal pathway**

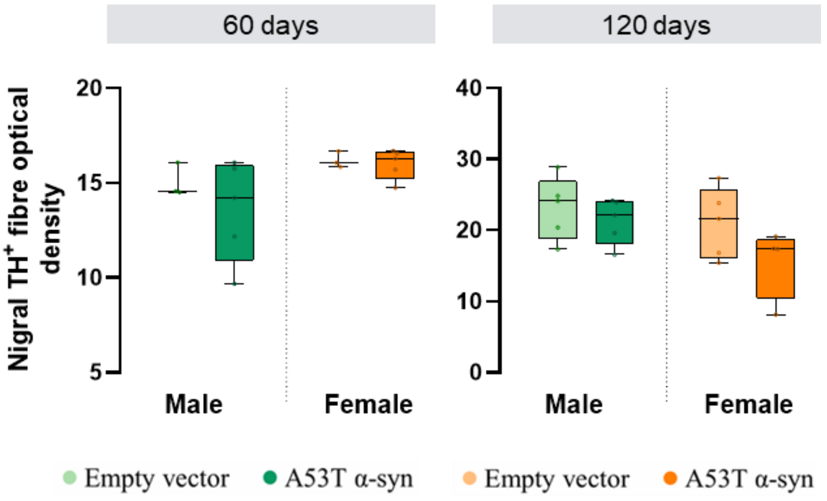


**Fig. S10. Quantification of the optical density of dopaminergic fibres in the *substantia nigra pars reticulata* (SNr) in male and female A53T alpha-synuclein (α-syn) and empty vector animals.** Dopaminergic neurons were positive for tyrosine hydroxylase ($TH^+$). No differences in dopaminergic degeneration were observed in the SNr at either 60 days or 120 days post-surgery.

**Supplementary Information References:**


Bidgood, R., Zubelzu, M., Ruiz-Ortega, J. A., & Morera-Herreras, T. (2024). Automated procedure to detect subtle motor alterations in the balance beam test in a mouse model of early Parkinson's disease. Scientific reports, 14(1), 862. https://doi.org/10.1038/s41598-024-51225-1

Brooks, S. P., & Dunnett, S. B. (2009). Tests to assess motor phenotype in mice: a user's guide. Nature Reviews Neuroscience, 10(7), 519-529. https://doi.org/10.1038/nrn2652

Goodwin, N. L., Choong, J. J., Hwang, S., Pitts, K., Bloom, L., Islam, A., ... & Golden, S. A. (2024). Simple Behavioral Analysis (SimBA) as a platform for explainable machine learning in behavioral neuroscience. Nature neuroscience, 27(7), 1411-1424. https://doi.org/10.1038/s41593-024-01649-9

Nath, T., Mathis, A., Chen, A. C., Patel, A., Bethge, M., & Mathis, M. W. (2019). Using DeepLabCut for 3D markerless pose estimation across species and behaviors. Nature protocols, 14(7), 2152-2176. https://doi.org/10.1038/s41596-019-0176-0
